# Toward understanding thrombus fracture: Dissipative phenomena of whole blood clots

**DOI:** 10.1101/2020.07.19.210765

**Authors:** Gabriella P. Sugerman, Sapun H. Parekh, Manuel K. Rausch

## Abstract

When thrombus fractures and breaks off it can occlude vital vessels such as those of the heart, lung, or brain. These thromboembolic conditions are responsible for 1 in 4 deaths world-wide. This problem is also of significant current interest as 1 in 3 COVID-19 intensive care patients exhibit thromboembolic complications. Thrombus resistance to fracture is driven by its intrinsic fracture toughness as well as other, non-surface-creating dissipative mechanisms. In our current work, we identify and quantify these latter mechanisms toward future studies that aim to delineate fracture from other forms of dissipation. To this end, we use an in vitro thrombus mimic system to produce whole blood clots and explore their dissipative mechanics under simple uniaxial extension, cyclic loading, and stress-relaxation. We found that whole blood clots exhibit Mullins effect, hysteresis, permanent set, strain-rate dependence, and nonlinear stress-relaxation. Interestingly, we found that performing these tests under dry or submerged conditions did not change our results. However, performing these tests under room temperature or body temperature conditions yielded differences. Overall, we have demonstrated that whole blood clots show several dissipative phenomena - similarly to hydrogels - that will be critical to our understanding of thrombus fracture.

## 1 Introduction

In vivo blood clots, or thrombi, play diametric roles in our body. Viscoelastic blood clots prevent hemorrhage after vascular injury and are thus vital to our well being ^1^. On the other hand, they are also the source of devastating thromboembolic conditions such as strokes, heart attacks, and pulmonary thromboembolism. In fact, 1 in 4 deaths world-wide are attributed to thromboembolic conditions ^2^. Thus, thromboembolic conditions are the number one cause of death. Additionally, 31% of patients under intensive care for COVID-19 have exhibited thromboembolic complications^3^.

Thrombus’s mechanical properties are important to both its physiological role as well as its pathological role. For instance, blood clots’ high extensibility and resistance to fracture are vital to their ability to seal injured vasculature and resist the pulsatile, circulatory load while the host tissue heals ^4^. On the other hand, it is when external forces overcome thrombus extensibility and resistance to fracture that they break off and embolization occurs, which precipitates the above mentioned thromboembolic conditions ^5^.

Blood clots’ mechanical properties result from the hierarchical interplay of their fibrin backbone, platelets, red blood cells, and pore-filling, interstitial plasma ^6,7^. During clot formation, fibrinogen is cleaved and activated to form a semi-flexible fibrin polymer ^8^. Fibrin is a remarkable polymer with extensibility up to 300% and significant fatigue resistance ^9–11^. Thus, it lends blood clots much of their high deformability. This backbone is stabilized and prestrained by activated or “sticky” platelets. These formed bodies attach to fibrin and actively pull on its fibers during the initial clot formation, thus, stiffening and densifying the material ^12,13^. Lastly, red blood cells contribute to clot mechanics by filling space and re-distributing loads. Although their role is passive, recent work has demonstrated that they drastically transform the behavior of their host material ^14^. Finally, driven by gradients in pore pressure, interstitial plasma permeates through the clot matrix and ostensibly contributes to its time-dependent behavior as it exchanges momentum with the surrounding matrix ^15^.

Because blood clots’ mechanical properties are critical in health and disease, they have been studied extensively. For a detailed review, we refer the reader to others’ excellent work ^16,17^. However, relatively few studies have focused specifically on thrombus properties that give rise to its ability (or inability) to resist external forces, i.e. embolize, with few exceptions ^18^. The study of thrombus fracture is complicated by its complex and time-variant composition and the resulting myriad non-linear mechanical phenomena contributing to its behavior ^19,20^. Generally speaking, resistance to fracture is driven by dissipative mechanisms ^21^. We categorize them as those that result in surface creation and those that are non-surface creating. In an effort to understand thrombus fracture as the biophysical phenomenon underlying thrombus embolization, we set out to characterize the non-surface creating dissipative mechanisms of whole blood clots. To this end, we employ an in vitro system in which we produce whole blood clots to mimic fresh thrombus. We tested these thrombus mimics with uniaxial tensile tests, cyclic tensile tests, and stress-relaxation tests to give a detailed account of the time-dependent mechanical properties that give rise to these non-surface-creating, dissipative mechanisms of thrombus.

## 2 Methods

### 2.1 Sample Preparation

We generated blood clots from bovine blood (Lampire Biological Laboratories, PA, USA) that was collected with CPDA-1 anticoagulant at 14% volume:volume and stored at 4°C for 12-72 hours prior to experimentation. To coagulate blood, we added calcium chloride to a final concentration of 20mM to reverse the anti-coagulant ^12,22^. To support coagulation, we gently mixed blood with a wide-mouth pipette before injecting it into a custom, 2-piece mold that was lined with hook-and-loop fabric on two sides, see Figure 1A. The mold, in turn, has sliding attachments for easy mounting to our table top Instron uniaxial tensile testing machine (Instron, Norwood, MA, USA). Thus, no direct clamping or gluing of the sample is necessary avoiding the danger of altering or damaging the material. We covered all samples during the 60 minute coagulation period to prevent dehydration ^23^.

**Fig. 1.**
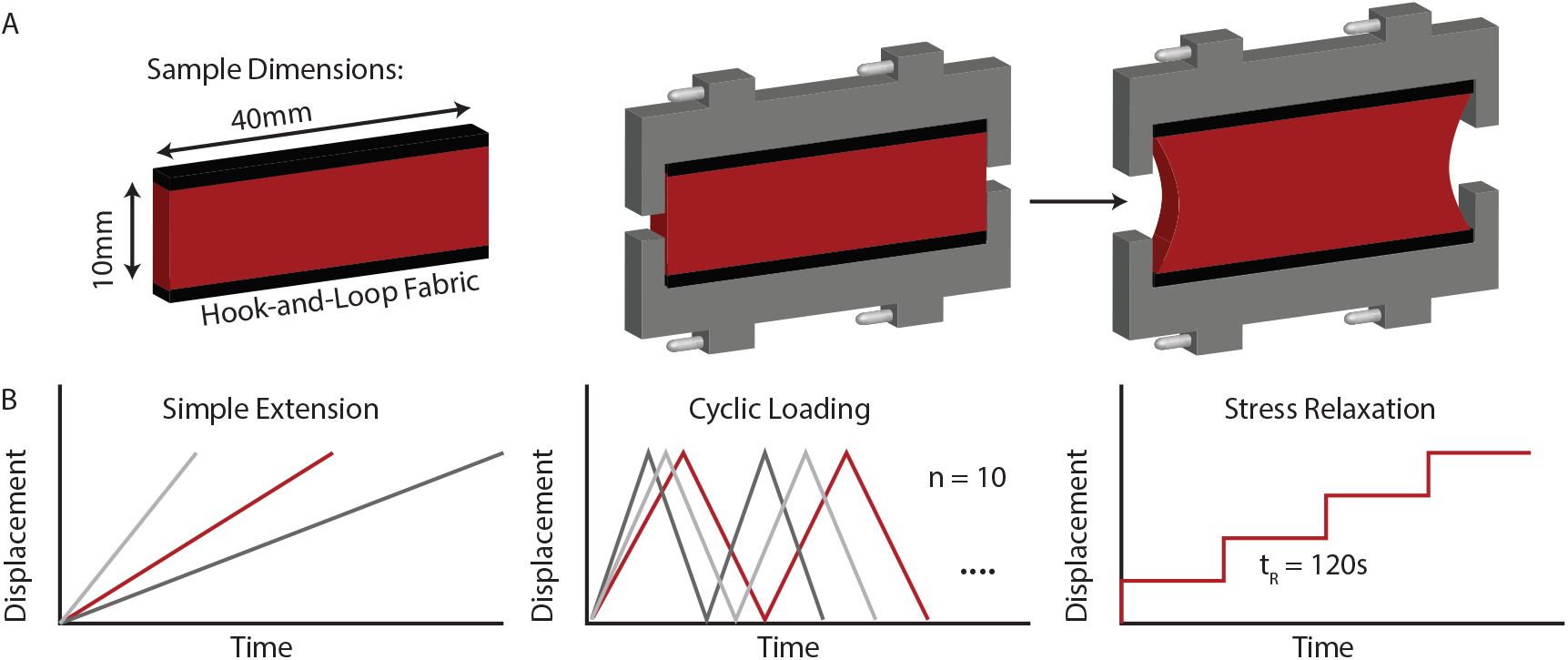
Sample dimensions and loading protocols. A) We coagulated whole blood into a custom, 2-piece mold that was lined with hook-and-loop fabric. B) During testing, we used an Instron to displace the upper mold piece according to three standard protocols: simple extension, cyclic testing, and stress relaxation tests. We conducted the simple extension and cyclic tests at three different strain rates

### 2.2 Mechanical Testing

After mounting the samples to the tensile testing machine, we displaced samples to 0.5mm to disengage the two-part molds. Subsequently, we conducted three types of uniaxial loading protocols: simple extension to 40% clamp-to-clamp strain, cyclic loading for ten cycles with a peak strain of 40%, and stress relaxation tests with four 120s holds at 10%, 20%, 30%, 40% strain. We conducted the simple extension and cyclic tests at three strain rates: 0.2mm/s, 1.0mm/s, and 5.0mm/s. During the stress-relaxation tests, we consecutively ramped the tissue to the four respective strains at a rate of 1mm/s. Additionally, we conducted all experiments under three different environmental conditions: at room temperature and without being submerged in phosphate buffered saline (PBS), being submerged in room temperature PBS, and being submerged in 37°C PBS. Finally, we prepared samples and conducted each experiment on Day 1, Day 2, and Day 3 after receipt of the blood.

### 2.3 Statistics

As described above, we designed a balanced study with each test type being performed for each condition on each day. Thus, we had one independent factor (blood batch, i.e., subject) and three dependent factors (strain rate, environmental condition, and storage time for the tensile and cyclic protocol, as well as strain, environmental condition, and storage time for the stress-relaxation protocol). We used the “afex” library as implemented in R to fit a linear mixed model to our data and the “emmeans” library to conduct multicomparisons between groups ^24–26^. All data are reported as mean curves with standard error curves. The sample numbers for each graph are provided in the figure captions.

## 3 Results

Our methodology of coagulating blood clots into hook-and-loop fabric-lined molds successfully yielded homogeneous, consistent, and well-secured samples. We tested these samples under uniaxial extension to answer fundamental questions about whole blood clots’ nonlinear, time-dependent mechanics.

### 3.1 Whole blood clots show a linear constitutive behavior and strain-rate dependence during initial loading

We first performed simple uniaxial tensile tests on unconditioned samples. Importantly, during pilot experiments, we tested samples to failure to determine the maximum strains before whole blood clots demonstrated signs of damage. Based on these tests, we evaluated the “elastic” behavior of whole blood clots for extensions up to 40%. Figure 2 shows extension curves for blood clot samples at 0.2, 1.0, and 5.0mm/s or equivalent strain rates of 2, 10, and 50%/s. These curves demonstrate that under uniaxial extension whole blood clots’ response was approximately linear. Furthermore, these samples’ stiffness depended on strain rate with their material response becoming increasingly stiff with increasing strain rate (p=0.016), i.e., whole blood clots mechanical behavior is strain-rate dependent.

**Fig. 2.**
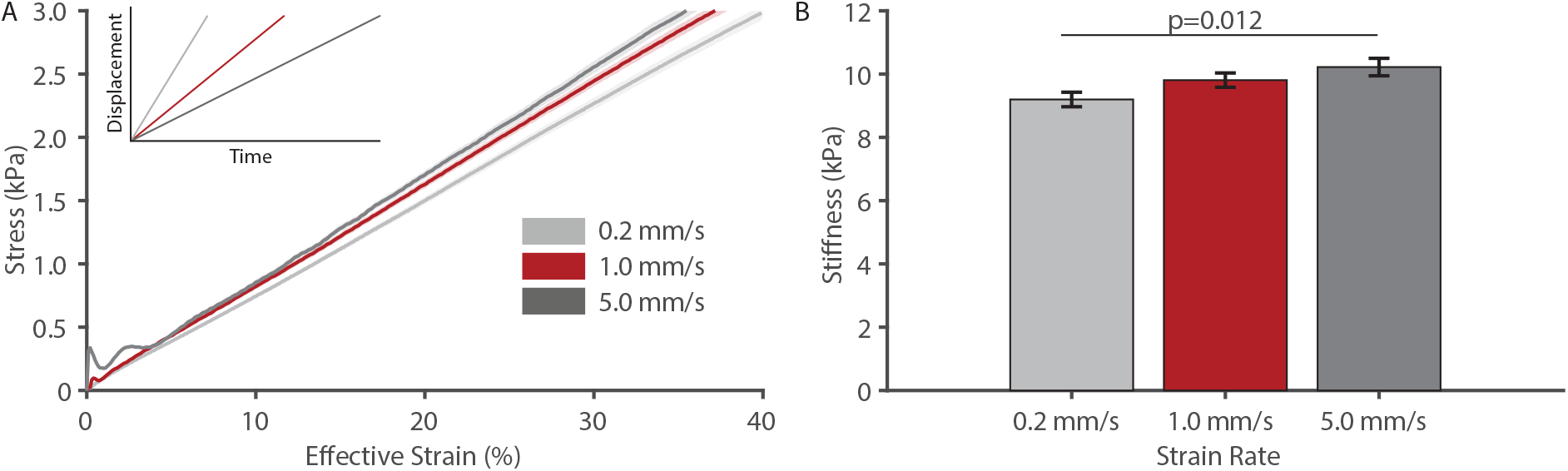
During initial loading whole blood clots behave linearly and strain-rate dependent. A) Stress-strain behavior of initially loaded whole blood clots at three strain rates (n=18 per group). B) Stiffness of whole blood clots at three different strain rates (n=18 per group). All data are mean ± standard error. Note, the initial spike in the 5.0mm/s curve is due to the motor acceleration and was not included for the stiffness calculation

### 3.2 Whole blood clots show hysteresis, Mullins effect, and permanent set

In Figure 3 we illustrate two distinct experiments to investigate the effect of whole blood clots’ history-dependence. First, we loaded samples to 40% strain, then unloaded them to their starting configuration at zero displacement, before reloading to 40% strain, see Figure 3A. In a second experiment, we unloaded the samples to zero force after the initial loading before returning to 40% strain, see Figure 3B. In the first experiment we observe that whole blood clots did not follow the initial loading curve during unloading. Instead, their response fell below the initial loading curve and entered a compressive state when approaching its reference configuration. This implies that i) whole blood clots dissipated energy during the initial loading cycle, and ii) that whole blood clots elongated during the original loading. The secondary loading to 40% strain followed neither the initial loading curve, nor the unloading curve. In other words, whole blood clots experienced Mullins in addition to hysteresis and permanent set. In the second experiment we observe the same qualitative and quantitative behavior. However, because we reversed the test direction once the samples reached a zero-force configuration (which is different from the reference configuration) we could estimate the permanent set accrued during the initial loading as approximately 12%. Additionally, comparison of both types of experiments at 0.2, 1.0, and 5.0mm/s reinforced our findings on blood clots’ strain rate-dependence.

**Fig. 3.**
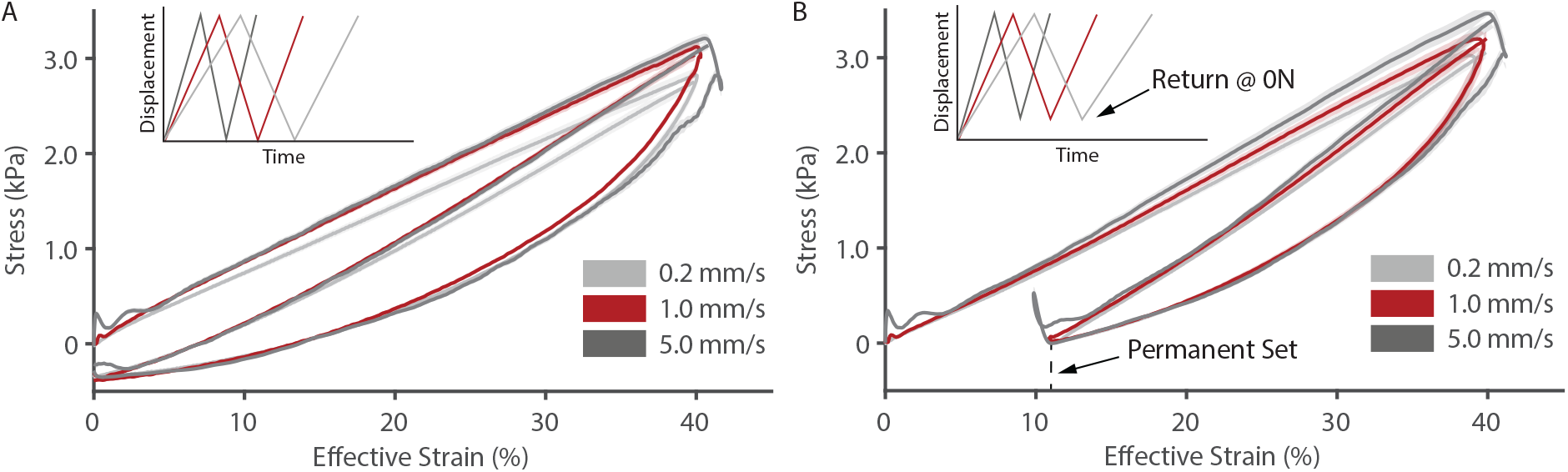
Whole blood clots show hysteresis, Mullins effect, and permanent set. A) A loading-unloading cycle with return to the original reference configuration (n=9 per group). B) A loading-unloading cycle with return to zero force, which allowed us to estimate permanent set to be approximately 12% (n=9 per group)

### 3.3 Whole blood clot mechanics equilibrate after ten cycles during which it transforms into a non-linear material

In vivo thrombus is exposed to cyclic forces due to shear and wall motion. Here, we performed cyclic loading in ten cycles to investigate whether whole blood clots approach an equilibrium state during those ten cycles. Specifically, we noted peak force, hysteresis, and permanent set for those ten cycles. In Figure 4A, we observed that blood clots showed preconditioning in that peak force, hysteresis, and permanent set dropped after the first cycles, but then trended toward an equilibrium response. While neither metric fully equilibrated by ten cycles, changes between the last and second-to-last cycle were small in comparison to the initial change between cycles one and two. Thus, ten cycles of loading may represent a practical compromise when simulating the equilibrated response of whole blood clots. Importantly, while whole blood clots show a linear stress-strain behavior during initial loading, they transform into a nonlinear material after ten loading cycles. Figure 4B compares the first and the tenth loading cycle illustrating the transformation from a linear to a nonlinear material. Note, for illustration purposes, we removed the permanent set after the tenth cycle.

**Fig. 4.**
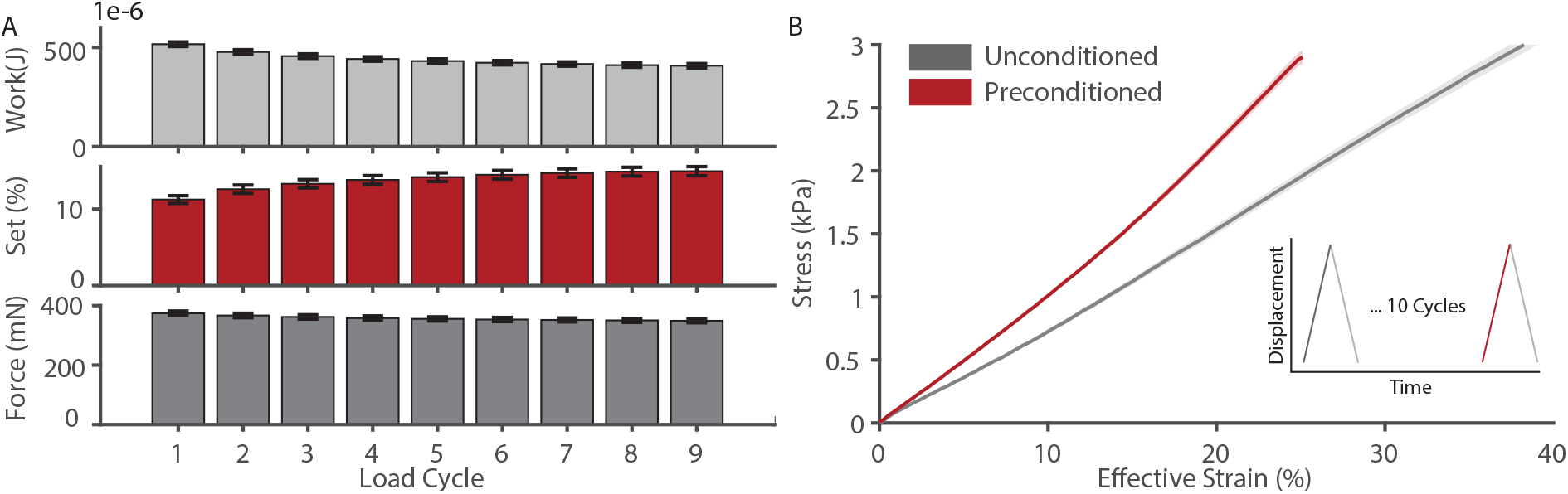
Whole blood clots’ mechanical response equilibrates after ten loading cycles. A) Work lost in hysteresis, permanent set, and peak force trended toward an equilibrium after ten loading cycles (n=9). B) After ten loading cycles, whole blood clots’ stiffness increased and took a slightly convex shape (n=9)

### 3.4 Whole blood clots undergo strain-dependent stress relaxation

To further investigate whole blood clots’ time-dependent behavior, we additionally performed sequential stress-relaxation experiments at 10%, 20%, 30%, and 40% strain, and let the material relax for 120s in between. We observed that blood clots relaxed at each strain, see Figure 5A. To quantify the relaxation behavior, we fit exponential decay functions to the data at each step, viz.

**Fig. 5.**
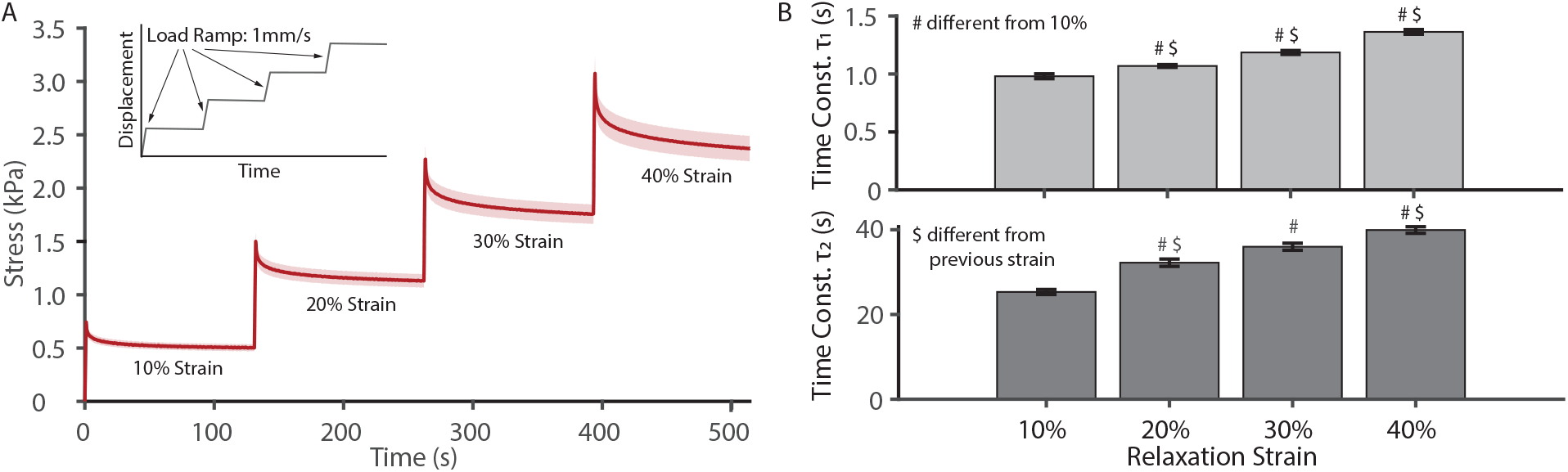
Whole blood clots are non-linear viscoelastic materials. A) Whole blood clots showed stress-relaxation (n=6). B) The relaxation response is well represented by an exponential decay function with two time constants both of which increased with strain (n=6)

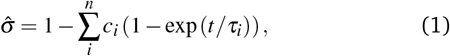

Where 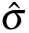 is the normalized stress at each given strain level, *t* is time, *c*_*i*_ are scaling parameters and *τ*_*i*_ are time constants. We compared the goodness of fit of a 1-term (n=1), 2-term (n=2), 3-term (n=3), and 4-term (n=4) function. We found that the goodness of fit measured as the normalized mean squared error (NMSE) significantly improved between one (NMSE = 0.79) and two terms (NMSE=0.98), but did not improve for three terms (NMSE = 0.98) or four terms (NMSE=0.98). Thus, we decided to report fits to a two-term function and identified two time constants, *τ*_1_ and *τ*_2_. Interestingly, our least squares fits consistently identified a short-term (around 1s) and a longer-term constant (around 30s), see Figure 5B. Both time constants increased with strain (p<0.001, p<0.001); thus, implying that whole blood clots show signs of nonlinear viscoelasticity similar to that reported by Nam et al ^27^.

### 3.5 Whole blood clots recover unloaded constitutive behavior after each relaxation step

When plotting stress against strain for the sequential stress-relaxation loading and comparing these curves to the unconditioned loading of the blood clot, we found that stress relaxation at lower strains does not affect clots’ constitutive behavior at larger strains, see Figure 6. Specifically, with increasing strain after each relaxation period stress returned to the unconditioned uniaxial loading behavior. This is only true when comparing the stress-relaxation data to the unconditioned loading curve at 1mm/s. When we compare the stress-strain behavior during the stress-relaxation tests against the unconditioned uniaxial behavior at 0.2mm/s, we find that it falls above the unconditioned uniaxial loading curve. These findings also imply that 0.2mm/s does not represent the quasistatic behavior of blood clot, which we predict would represent the lower bound of the stress-strain curve during stress-relaxation.

**Fig. 6.**
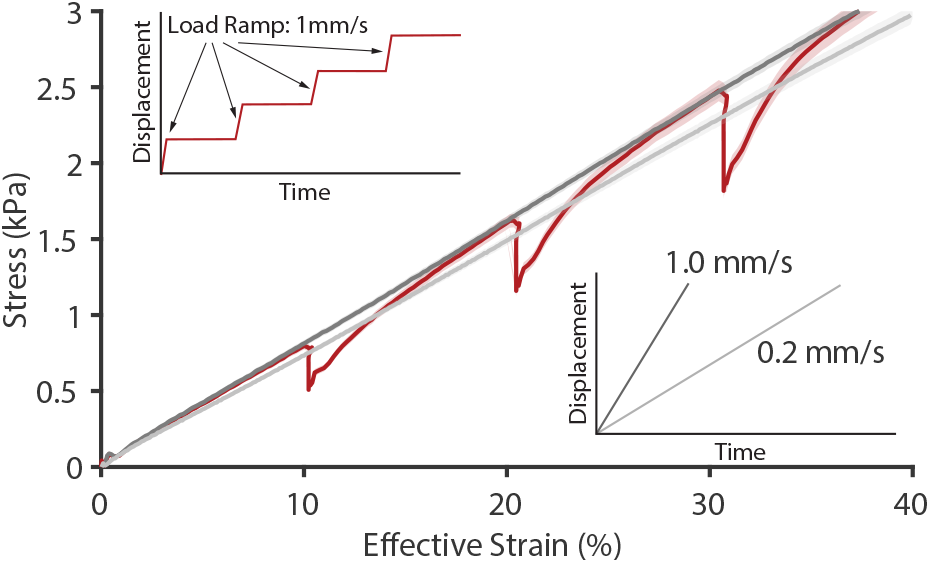
Whole blood clots recover unloaded constitutive behavior after each stress-relaxation test. We show the stress-stress behavior extracted from the sequential stress-relaxation tests (red, as shown in Figure 5A, n=18) and compared it to the unloaded behavior under simple extension at both 0.2mm/s (light gray, n=18) and 1.0mm/s (dark gray, n=18) (as shown in Figure 2A)

### 3.6 Whole blood clots’ stress-relaxation not strongly dependent on geometry

The identification of two time constants may imply that two distinct physical processes contribute to stress relaxation. Given the highly hydrated state of whole blood clots and the polymeric nature of blood clots’ fibrous backbone, it is possible that diffusion-based and solid viscoelasticity-based processes may collaboratively determine blood clots time-dependent behavior. While the former would be expected to be a function of sample cross-section, the latter would not. To investigate the degree to which these time constants and thus the underlying physical processes depend on geometry, we repeated above experiments on samples of 4mm and 5mm cross-section (as opposed to all other samples that had 3mm cross-sections). Table 1 shows a comparison between the time constants between those three sample types. Neither the short-term response nor the longer-term response appeared to significantly increase with increasing cross-section (p=0.287 and p=0.289). This may support the hypothesis that time-dependence resulted from solid viscoelasticity rather than diffusion-based processes. However, future studies with larger geometric range may be able to shed a more confident light on this question.

**Table 1.** Time constants *τ*_1_ and *τ*_2_ as a function of sample size and strain. Data shown as mean ± standard error (n=6 per group)

| $\tau_1$ | | | | |
| --- | --- | --- | --- | --- |
| Size | 10% | 20% | 30% | 40% |
| 3mm | $1.09 \pm 0.02$ | $1.18 \pm 0.04$ | $1.31 \pm 0.04$ | $1.42 \pm 0.02$ |
| 4mm | $1.08 \pm 0.01$ | $1.18 \pm 0.01$ | $1.30 \pm 0.01$ | $1.41 \pm 0.01$ |
| 5mm | $0.97 \pm 0.03$ | $1.14 \pm 0.02$ | $1.26 \pm 0.01$ | $1.43 \pm 0.02$ |
| $\tau_2$ | | | | |
| Size | 10% | 20% | 30% | 40% |
| 3mm | $28.7 \pm 0.6$ | $29.9 \pm 1.4$ | $32.4 \pm 0.6$ | $33.6 \pm 0.2$ |
| 4mm | $25.7 \pm 0.9$ | $29.5 \pm 0.3$ | $31.2 \pm 0.3$ | $33.2 \pm 0.5$ |
| 5mm | $23.6 \pm 0.6$ | $28.2 \pm 0.4$ | $31.3 \pm 0.3$ | $33.5 \pm 0.2$ |

### 3.7 Whole blood clot mechanics do not depend on humidity conditions, but depend on temperature

Given that thrombi form within vessels, their natural environment is within a physiological fluid bath and at body temperature. We have previously tested whole blood clots only under those conditions. As testing blood clots under non submerged, room-temperature conditions would ease experimental protocols, we wanted to test the effect of both temperature and humidity on the mechanics of whole blood clots. Thus, we compared the mechanics of blood clots when tested at room temperature without a bath, in a room-temperature PBS bath, and in a 37°C PBS bath, both under simple extension and during stress-relaxation. Figure 7A shows the results for the simple uniaxial extension tests, while Figure 7B shows the results for the stress-relaxation tests as a function of those conditions. In the former figure, variations appear small suggesting that stiffness was not affected by testing conditions (p=0.717). However, differences seem to exist in the stress-relaxation data. A closer look revealed that the constant *τ*_1_ was shorter in the 37°C bath than under the other two conditions (p<0.001). Interestingly, we did not find that *τ*_2_ depended on temperature (p=0.680).

**Fig. 7.**
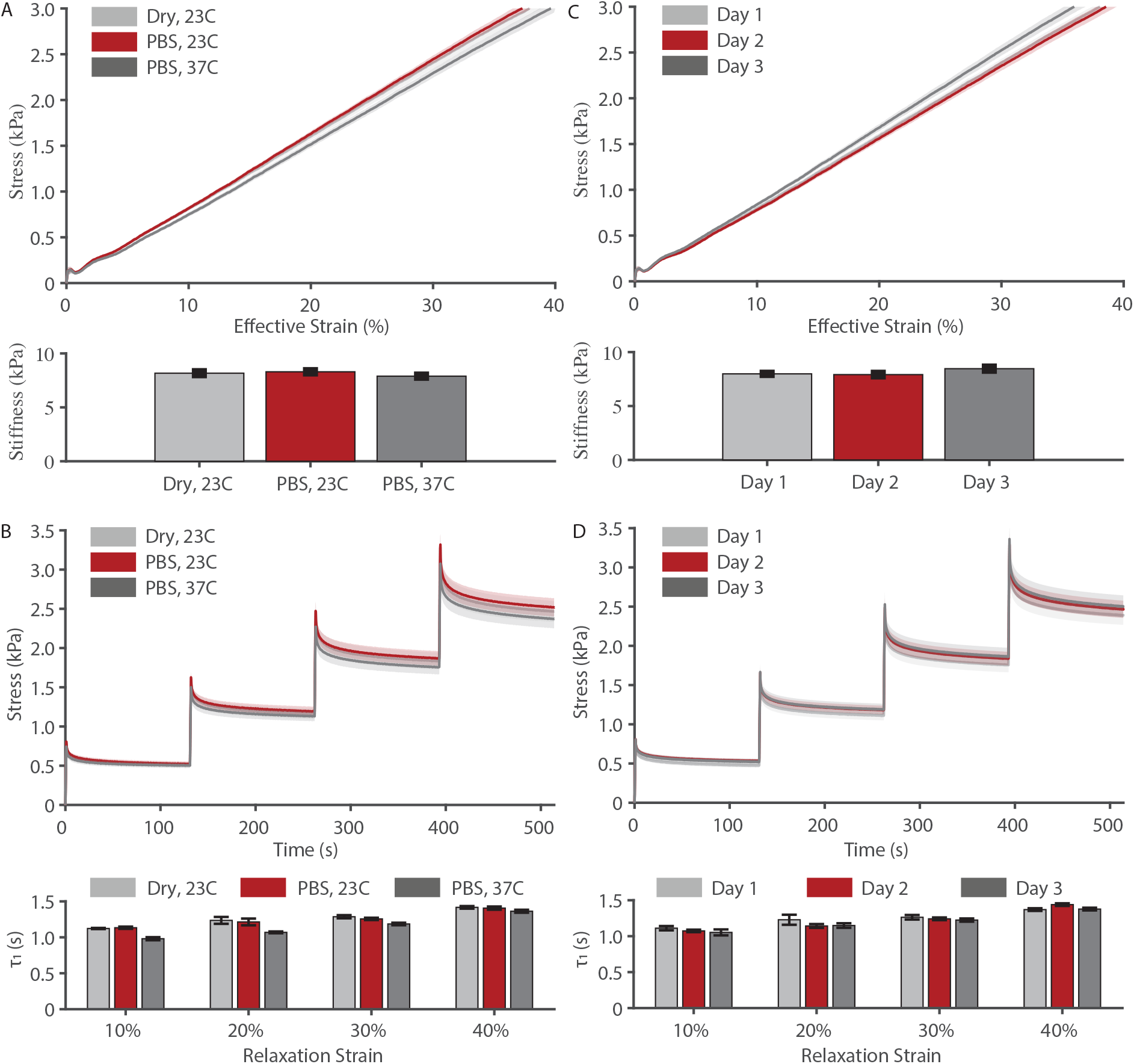
Whole blood clots’ non-linear, time-dependent behavior depends on test temperature, but not hydration conditions during testing or on blood storage time. A) Simple uniaxial extension of whole blood clots as a function of environmental conditions (n=18 per group). B) Stress-relaxation of whole blood clots as a function of environmental conditions (n=18 per group). C) Simple uniaxial extension of whole blood clots as a function of blood storage time (n=6 per group). D) Stress-relaxation of whole blood clots as a function of blood storage time (n=6 per group)

### 3.8 Whole blood clots’ relaxation behavior, but not its stiffness, depends on storage time

Similarly to our test of whole blood clots’ mechanical property dependence on environmental conditions, we also wanted to investigate the effect of blood storage time. While we have previously performed a similar analysis under simple shear, during which we found that whole blood clots’ stiffness did not depend on blood storage time during days one through three, we only did so under one loading rate and did not consider the time-dependent behavior ^23^. Figure 7C and 7D show whole blood clots’ uniaxial loading behavior and stress-relaxation behavior as a function of blood age, comparing one day, two days, and three days of blood storage before coagulation and testing. We found slight differences between day one and three in the first time constant, which were not statistically significant *τ*_1_ (p=0.068). We suspect that increased sample number would provide the necessary power to statistically support this difference. However, we did not see any differences in whole blood clots’ stiffness, which is in agreement with our previous findings (p=0.121) ^23^.

## 4 Discussion

While blood clot and thrombus mechanics have been extensively investigated, comprehensive mechanical studies focusing on their time-dependent properties have been sparse; moreover, studies investigating their fracture behavior are all but absent. This gap should be overcome as thrombus fracture leads to thromboembolic conditions, which is a substantial national and global health problem. Generally speaking, fracture occurs when external work on a body produces a sufficient stress concentration and supplies sufficient energy to create new surfaces ^28^. The material’s intrinsic ability to resist the creation of new surfaces may be quantified in its fracture toughness, a material property ^29^. However, energy dissipating mechanisms, such as those studied here, may lower the energy available for surface creation, thus, increasing the apparent toughness of the material ^30^. This non-surface-creating dissipation of energy is the basis for toughening strategies of both classic engineering materials such as steel as well as non-classic engineering materials such as hydrogels ^31,32^. For example, in double network hydrogels, one network family, the sacrificial network, yields at relatively small loads and thereby dissipates energy that protects the non-sacrificial network from fracture ^33^.

Before endeavoring on identifying the surface-creating dissipative mechanisms during whole blood clots’ fracture, we felt it was prudent to first identify the non-surface-creating dissipative mechanisms. We anticipated that we’d observe damage, hysteresis, stress-relaxation, and strain-rate dependence ^34^. Each of these phenomena contributes (or is an example of) energy dissipation before fracture. It is only after identifying and quantifying these phenomena that we can deconvolute the mechanisms that lead to thrombus ability (or inability) to resist external forces. Additionally, we tested the effect of storage and environmental conditions on these mechanisms for future reference.

As we expected, we observed that whole blood clots were strain-rate dependent. Albeit, across the 25-fold increase in strain rate, the materials stiffness increased only marginally. Additionally, we found that whole blood clots dissipate energy during the first loading-unloading cycle via Mullins effect ^35^, which is possibly enhanced compared to pure fibrin by the inclusion of RBCs as classical filler particles ^14,36^. This initial energy dissipation due to the Mullins effect is compounded through hysteresis, which we continued to observe during subsequent loading cycles, not just the first. Interestingly, we found that whole blood clots’ response to cyclic loading equilibrated at around ten cycles. Thus, repeated or cyclic loading as seen in vivo likely alters thrombus ability to resist fracture only during the first cardiac cycles and stabilizes afterwards. Note, that fatigue, which we have not investigated here, may weaken the material over time at cycle numbers larger than ten ^37^. We also found that whole blood clots relax under strain. This response changes with increasing strain indicating that whole blood clots behave like a non-linear viscoelastic material ^27,38^.

In our stress-relaxation experiments, we found two relaxation timescales, possibly suggesting a diffusion-based and solid viscoelasticity-based relaxation. With our additional experiments showing geometry-independence of the relaxation constants, our data indicate that the time-dependent behavior of whole blood clots may be driven by solid viscoelasticity, rather than diffusion-based mechanisms. Alternatively, the two time scales could be founded in processes on different scales, e.g., network-scale, fiber-scale, protofibrillar-scale, or molecular-scale. For instance, strain stiffening of fibrin gels is driven by such hierachical phenomena ^39^.

In our work, we also found that whole blood clots’ nonlinear, time-dependent behavior was insensitive to whether tests were conducted in a fluid bath or under dry conditions. This finding may additionally support the hypothesis that blood clots’ time-dependent mechanical properties are driven by solid viscoelasticity rather than diffusion-based viscoelasticity. However, we did find that these properties were slightly different when conducted at room temperature or at body temperature. Similarly, we found marginal effects of blood storage conditions on these properties. Dependence on storage conditions is likely driven by death of red blood cells and platelets, which we found to occur within days of storage ^23^. Our data provide an incentive to test whole blood clots and thrombus at body temperature and with short blood storage. If future studies should ignore these recommendations, our data provide the means to estimate the error accrued by doing so.

We propose that future studies on the fracture of thrombus - or whole blood clots as thrombus mimics - may follow an approach as outlined by Zhang et al in their study of hydrogel fracture ^30^. Specifically, they combined a nonlinear finite element model with mode I fracture experiments to delineate a hydrogel’s intrinsic and effective toughness. Our current work provides the information and data to do so. Additionally, we are currently lacking an established constitutive model that may cast our data into mathematical and numerically-amiable form. Future studies may also want to explore the best constitutive frameworks and specific forms.

## 5 Conclusions

Whole blood clots show a number of non-surface-creating, dissipative phenomena prior to fracture. These materials show clear evidence of Mullins effect, hysteresis, permanent set, and non-linear stress-relaxation. Quantifying these phenomena is critical to our understanding of thrombus fracture as the biophysical phenomenon underlying embolization. With support of our work and our data, future studies will be able to focus on explicit fracture, i.e., the creation of surfaces via crack propagation while accounting for other dissipation behavior.

## Conflicts of interest

Manuel K. Rausch has a speaking agreement with Edwards Life-sciences. None of the other authors have conflicts to declare.

## Acknowledgements

This work was partially funded by the K.C. William’s Faculty Excellence Fund through the University of Texas at Austin Aerospace Engineering & Engineering Mechanics Department. The authors are also very appreciative for frequent and helpful discussions with Emma Lejeune.

## Notes

### Competing Interest Statement

The authors have declared no competing interest.

